# Dynamic phenotypic heterogeneity generated by delayed genetic oscillations

**DOI:** 10.1101/2020.05.13.093831

**Authors:** R. Peña-Miller, M. Arnoldini, M. Ackermann, R. E. Beardmore

## Abstract

Eukaryotes and prokaryotes exploit the ability of genetically identical cells to exhibit different phenotypes in order to enhance their survival. However, the mechanisms by which cells transition from one phenotype to another remain unclear. Canonical models of this dynamic posit that molecular fluctuations provide the noise that drives the cell out of one stable state and into another. Stochastic processes generated by canonical models should, therefore, be good descriptors of phenotype dynamics and between-state transitions should become more likely at greater noise amplitude, for instance at higher extracellular temperatures. To test these predictions, we observed temporal expression dynamics of the promoter of a flagellum gene, *fliC*, in a microfluidic device using *Salmonella enterica* serovar Typhimurium and green fluorescent protein (GFP). Our observations show that while cells can exhibit multistable phenotypes, including stable *fliC*-OFF and *fliC*-ON states characterised by low and high GFP levels, respectively, between-state transitions can exhibit oscillatory dynamics whose return statistics do not conform to canonical theories. For example, here the *fliC*-ON state was more frequent following a temperature increase. To better understand our data we developed different dynamical frameworks to predict *fliC* expression data. We conclude that a stochastic dynamical system tailored to the genetic network of *fliC* is better suited to our data than prior theories where dynamical features, like oscillations and pulsing, are driven by inevitable delays in the post-translational regulation of *fliC*. Thus, while transcriptional noise promotes phenotypic heterogeneity, as we show here, regular features like oscillatory heterogeneity can result from delays that fundamental molecular processes impose upon a cell’s gene regulatory architecture.

## Introduction

Phenotypic heterogeneity – the expression of different phenotypes by clonal individuals living in a homogeneous environment – is pervasive among living cells. It is not merely the consequence of a cell’s inability to tightly regulate gene expression, but phenotypic heterogeneities can have adaptive functions.^1^ These can be broadly organised into two groups. First, individuals that express different phenotypes can interact with each other and divide labor. Such division of labor has been observed in the context of bacterial infections where individuals that express different sets of genes specialise on different roles in the infection process.^2^ It has also been observed in the context of microbial metabolism where individuals in clonal populations sometimes specialise on different metabolic pathways that complement each other.^1, 3–6^ Second, phenotypic heterogeneity can allow a genotype to persist in the face of environmental fluctuations. Phenotypic variation between individuals in clonal populations increases the likelihood that at least some individuals express a phenotype that allows them to survive and grow when external conditions suddenly change. This phenomenon, known as bet-hedging, has been observed in contexts including sporulation,^7^ bacterial persistence during exposure to antibiotics^8^ and metabolism.^9, 10^ Bet-hedging strategies can be selected for when sensing external cues and regulating metabolic machinery accordingly provides limited protection in unpredictable environments,^11^ or if the timescale of metabolic regulation is larger than the frequency of the fluctuations.^12^ In such cases it can be optimal for a population to be composed of multiple sub-populations at all times, each expressing a different phenotype.^13^

Phenotypic heterogeneity, and more specifically the stable co-existence of multiple phenotypes in a clonal population, is generally assumed to be a consequence of the inherent stochasticity of gene expression.^14^ Indeed, noise is difficult to avoid inside a cell: transcription and translation can be seen as a series of biochemical reactions characterised by molecular kinetics that can involve a small number of molecules^15^ and, therefore, phenotypic noise can arise as a consequence of temporal fluctuations in mRNA and protein numbers.^16, 17^ Specific features in the underlying genetic architecture, such as feedback loops, can translate these fluctuations into discrete phenotypes, so that individuals with the same genotype can exists in two or more distinct phenotypic states. Previous studies have focused on identifying the sources of stochasticity, for instance the amplification of gene expression noise,^18, 19^ fluctuations in the numbers of molecules present at low concentrations,^20^ random binding kinetics of promoters,^21^ stochastic partitioning during cell division^22^ and the amplification of noisy responses to biochemical signals in metabolic pathways.^23^

Perhaps the simplest noise-driven mechanism that produces phenotypic heterogeneity is the bistable toggle-switch. This mechanism generates a bimodal distribution of phenotypes by letting each cell in a population stochastically switch the expression of a noisy target gene between ON and OFF states. Such bimodal phenotypic distributions have been observed in a number of experimental systems, for example, in the induction of the *lac*^24^ and *ara*^25^ operons, as well as the bacteriophage λ lysis-lysogeny decision circuit.^26^ Furthermore, a synthetic toggle-switch was constructed based on two mutually repressive negative feedbacks and shown to achieve robust bistable behaviour experimentally.^27^ But is phenotypic bistability exclusively driven by stochastic fluctuations? Or could we view the cell as a dynamical system, indeed almost deterministically, but one subject to stochastic forcing, that regulates and adapts aspects of its phenotype dynamically through time?

Here we answer this question by showing, using mathematical models, computational simulations and single-cell microfluidics, that bistable toggle-switches can exhibit complex oscillatory dynamics, thus ensuring that each cell displays different phenotypes at different times. We focus on the expression dynamics of genes associated with the synthesis of a flagellum in a pathogenic strain of *Salmonella enterica* serovar Typhimurium (hereafter referred to as *Salmonella*), a virulence factor regulated by a gene network with an architecture characteristic of a bistable toggle-switch: a double-negative feedback loop.^28^ Moreover, the expression of these genes is known to be heterogeneous in *Salmonella* populations.^29–31^

Phenotypic heterogeneity in the expression of flagellar genes is functionally relevant during the growth of *Salmonella* in a host. The expression of flagella allows these bacteria to move in their environments, enabling them to find nutrient sources and host cells to infect, but motility comes at a cost. Flagella are important antigens recognised by the mammalian immune system, triggering inflammation and the generation of antibodies tailored to seek and kill infecting bacteria.^32, 33^ *Salmonella* can mitigate this cost by switching between one of two types of flagella using a process known as flagellar phase variation,^34^ and also by creating a phenotypic sub-population that does not express flagellar proteins.^28, 29, 32, 35^ Heterogeneity in the expression of flagella has been suggested to have functional consequences aside from immune evasion, for instance by allowing the invasion of host cells when combined with the expression of the *Salmonella* Pathogenicity Island 1 gene cluster (also heterogeneously expressed in *Salmonella*^2, 31, 36^).

We show that the gene regulatory network controlling flagellar gene expression is consistent with a bistable toggle-switch model whereby phenotypic transitions are driven by a stochastic process. However, experimental time-series of gene expression in single cells reveal different gene expression dynamics to those predicted by the toggle-switch model. To explain this discrepancy mechanistically, we use mathematical models and numerical simulations to recapitulate single-cell data. We argue that the temporal dynamics of gene expression in single cells can be explained by a deterministic signal in the midst of the noise: delay-induced genetic oscillations. Our results show that phenotypic heterogeneity can result from deterministic dynamics hard-wired into the architecture of genetic networks, potentially allowing for enhanced controllability in the presence of noise.

## Results

### Gene regulatory dynamics of *fliC* expression

The gene regulatory network underlying heterogeneity in flagellar gene expression has been studied extensively.^37–40^ At its core lies a double negative feedback loop, which makes its architecture consistent with a bistable toggle switch. In short, the transcriptional master regulator FlhDC binds to promoters that increase transcription of operons encoding for structural proteins of the flagellar base and other regulatory proteins, including FliA (an alternative sigma factor) and FliZ. FliA specifically induces the expression of further flagellar proteins, including FliC, the structural protein that forms the flagellar filament. More recently, YdiV was reported to repress the master regulator FlhDC^41^ by post-translationally interacting with the FlhD subunit. *ydiV* transcription is in turn repressed by FliZ,^42^ thus producing a double negative feedback loop architecture: YdiV inhibits *fliAZ* expression by encoding an anti-FlhDC factor, while FliZ represses the transcription of *ydiV* (see Figure 1A for a graphical representation of this pathway).

**Figure 1.**
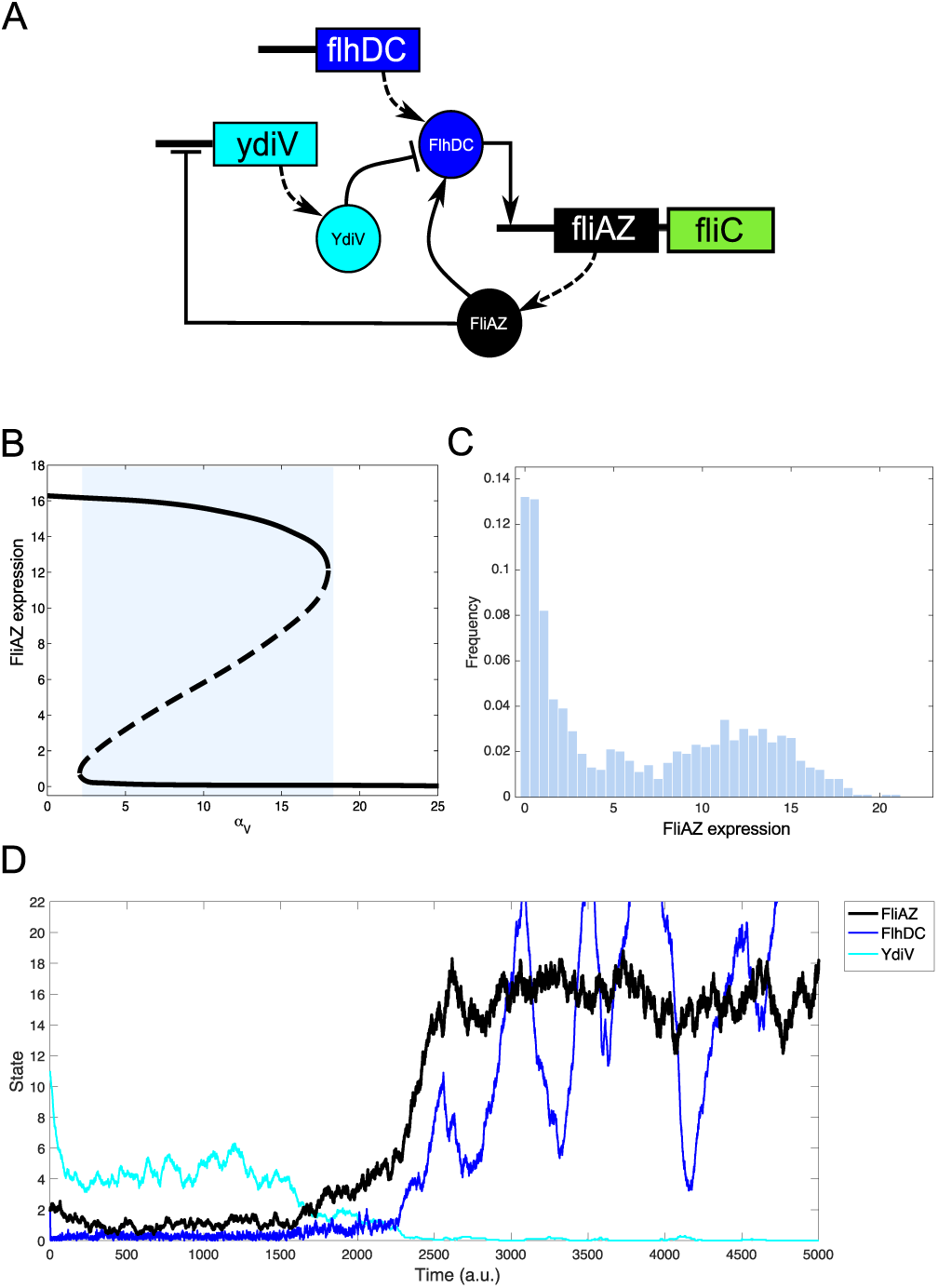
A) Network diagram representing the interaction between the three-tier operon regulating the expression of the flagellum. B) Bifurcation phase diagram of a gene regulatory model showing that the system presents bistability for certain parameter values (in this case, *α*_*V*_, the rate of interaction between *YdiV* and *FlhDC*). Solid lines represent stable steady states, the dotted is an unstable branch and the shaded area the parameter region where the system presents bistability. C) Theoretical frequency distribution of the expression of *fliC* simulated using the stochastic model with 1000 cells. As expected in a bistable toggle-switch, the resulting histogram of gene expression presents bimodality. D) Stochastic simulation of the bistable toggle-switch model (black solid line represents the expression level of FliAZ, Ω = 0.05). Note how at *t* = 0 the state of the system is in the *fliC*-OFF state, until transcriptional noise drives the system towards the *fliC*-ON steady state. Genes are denoted with rectangles and proteins with circles.

In order to understand the gene regulatory dynamics that such a system would encode, we modelled the dynamics of the flagellar regulon based on these regulatory interactions. We assume that the concentrations of proteins and mRNAs can be represented by non-negative, continuous time-dependent variables and that positive and negative regulatory interactions can be described by a non-linear Hill function. Therefore, we modelled the dynamics of flagellar regulation using a system of differential equations (equations detailed in the Methods section) and used numerical tools to study the dynamics of that system.

As illustrated by the phase diagram in Figure 1B, there exists a range of parameter values where the system exhibits bistability. These numerical results are consistent with prior studies showing that FlhDC acts as a master regulator of the flagellar toggle-switch.^30, 41^ Indeed, when FlhDC is expressed, the transcription of flagellar proteins is promoted and the dynamics of the system converges to the *fliC-ON* state. However, if YdiV is expressed, it suppresses transcription of *fliAZ* by sequestering FlhDC and hence the system is driven to the *fliC-OFF* state. Figure S1 shows the results of numerical simulations with the same parameter values but a different initial *ydiV* concentration; as a result, the system converges towards a different state. This dependence of final outcome on initial conditions suggests that transitions between basins of attraction of the phenotypic states can be driven by stochastic variations in protein or mRNA concentrations.^43^

We use a computational approach that simulates a gene regulatory model posed as a system of differential equations using a standard Monte Carlo algorithm proposed by Gillespie^44^ (see Methods and Table S1). This approach is used widely to model stochastic gene regulatory dynamics^45, 46^ because it provides a systematic method for obtaining a trajectory ensemble with statistics that asymptotically converge to the solution of the corresponding master equation (see Figure 1D for a realisation of the stochastic model). Moreover, this simulation-based approach allows us to produce theoretical population-level histograms of gene expression. For example, Figure 1C shows the frequency distribution of FliAZ expression of 100 cells with identical initial conditions and kinetic parameters (described in Table S2). Interestingly, the bimodal distribution obtained from our synthetic data is analogous to the experimental distribution obtained by flow cytometry (Figure S2).

Given the correspondence between our population-level experimental data and the numerical simulations, it would seem natural to conclude that a bistable toggle-switch model accurately describes the regulatory dynamics underlying phenotypic heterogeneity in flagellar gene expression. However, the observed phenotype distributions can also be described by different underlying dynamics at the single-cell level. For instance, it has been reported that high-levels of noise in a graded switch with a sigmoidal response can also produce bimodal distributions in gene expression data,^47^ so too can complex non-linear dynamics, notably oscillatory behaviour.^48, 49^ To avoid making inferences about the underlying gene regulatory network that produces the observed population-level distribution of gene expression, we use time-resolved gene expression data taken from individual cells grown in microfluidic devices observed under a microscope.

### Single-cell time-series of *fliC* expression

In recent years the development of microfluidic technology has allowed the study of bacterial growth and gene expression at a single-cell level. The utility of such devices derives from their ability to control the extracellular environment and to follow the life history of individual cells, for instance by measuring elongation and division rates or quantifying gene expression dynamics using fluorescent proteins. We used a microfluidic device commonly known as a *mother-machine*^50^ to obtain single-cell profiles of gene expression of 678 individual cells with a low copy number plasmid containing the gene *gfpmut2* encoding a green fluorescent protein under control of the *fliC* promoter. Time-lapse movies were obtained by acquiring images every 5 minutes for approximately 700 minutes. Using bespoke image processing algorithms we obtained time-series of fluorescence intensity for each cell, and normalised it with respect to population-level mean fluorescent intensity (Figure 2A shows a temporal montage of still images from different time points).

**Figure 2.**
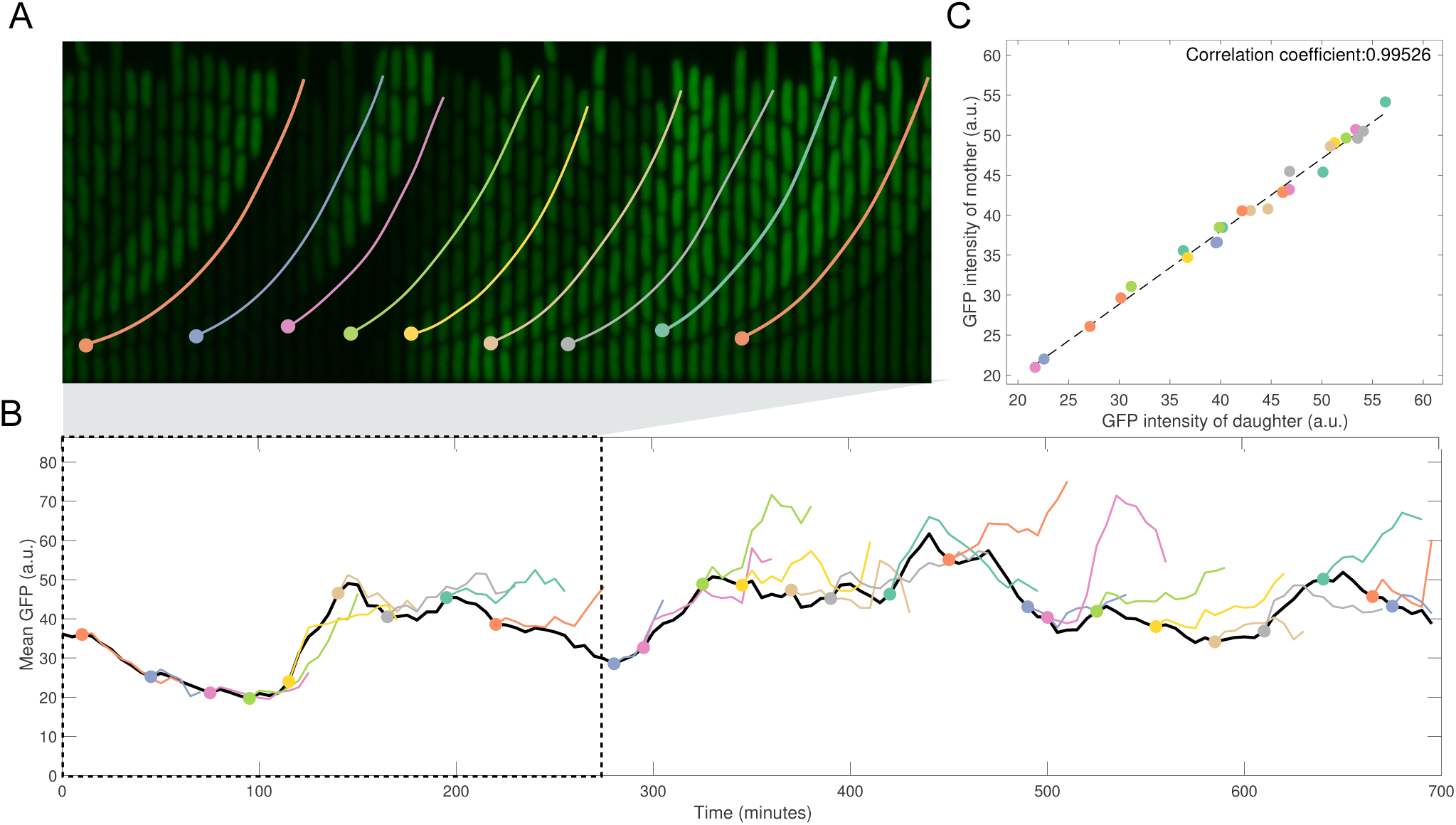
A) Temporal montage obtained from a single microfluidic channel illustrating how we can observe division events and quantify time-resolved data of GFP fluorescent intensity of a focal cell. Colour lines represent the tracking of the daughter cells until they are washed out of the microfluidic chamber. B) Fluorescent intensity of the mother and daughter cells are correlated at the moment of division, suggesting that the dynamics of gene expression is not coupled to the cell division cycle. C) Mean GFP as a function of time for a focal cell (black line). Circles represent division events and coloured lines the fluorescent intensity of the daughter cell (coloured lines are shorter, as daughter cells eventually are removed from the microfluidic channel).

To exploit our time-resolved data, we proceeded to analyse the temporal dynamics of gene expression. Figure 2 shows the expression profiles of a mother cell and her daughters before they are washed out of the channel. (The cell with the oldest cell pole resides at the bottom of each channel, and we refer to this cell as the mother cell. The daughters are young-pole cells that emerge when the mother cell divides. The daughter cells are located above the mother cell in the channel and are eventually removed from the microfluidic device). Notably, the expression profile of the daughter and mother cells are correlated at the moment of division (Figure 2C), suggesting that transitions between different states are not driven by stochastic partitioning during cell division. Moreover, the GFP expression profile of the daughter cells seems to follow the dynamics of the mother cell until both trajectories eventually diverge (Figure 2B).

As predicted by the toggle-switch model, a subset of cells are not expressing fluorescence for the entire duration of the experiment and therefore the state of the network is always in the *fliC*-OFF steady state in those cells see (Figure 3A-B). Conversely, cells expressing fliC exhibit complex temporal patterns that fluctuate between *fliC*-ON and *fliC*-OFF states (Figure 3A-B). We used this dataset to estimate the expression profile of *fliC* (as measured by GFP intensity) and estimated a population-level histogram of GFP expression. As Figure 3C shows, the latter is a bimodal distribution of GFP intensity as predicted by the toggle-switch model (Figure 1C); these data from single-cell microscopy experiments are consistent with the GFP distributions obtained using flow cytometry (Figure S2).

**Figure 3.**
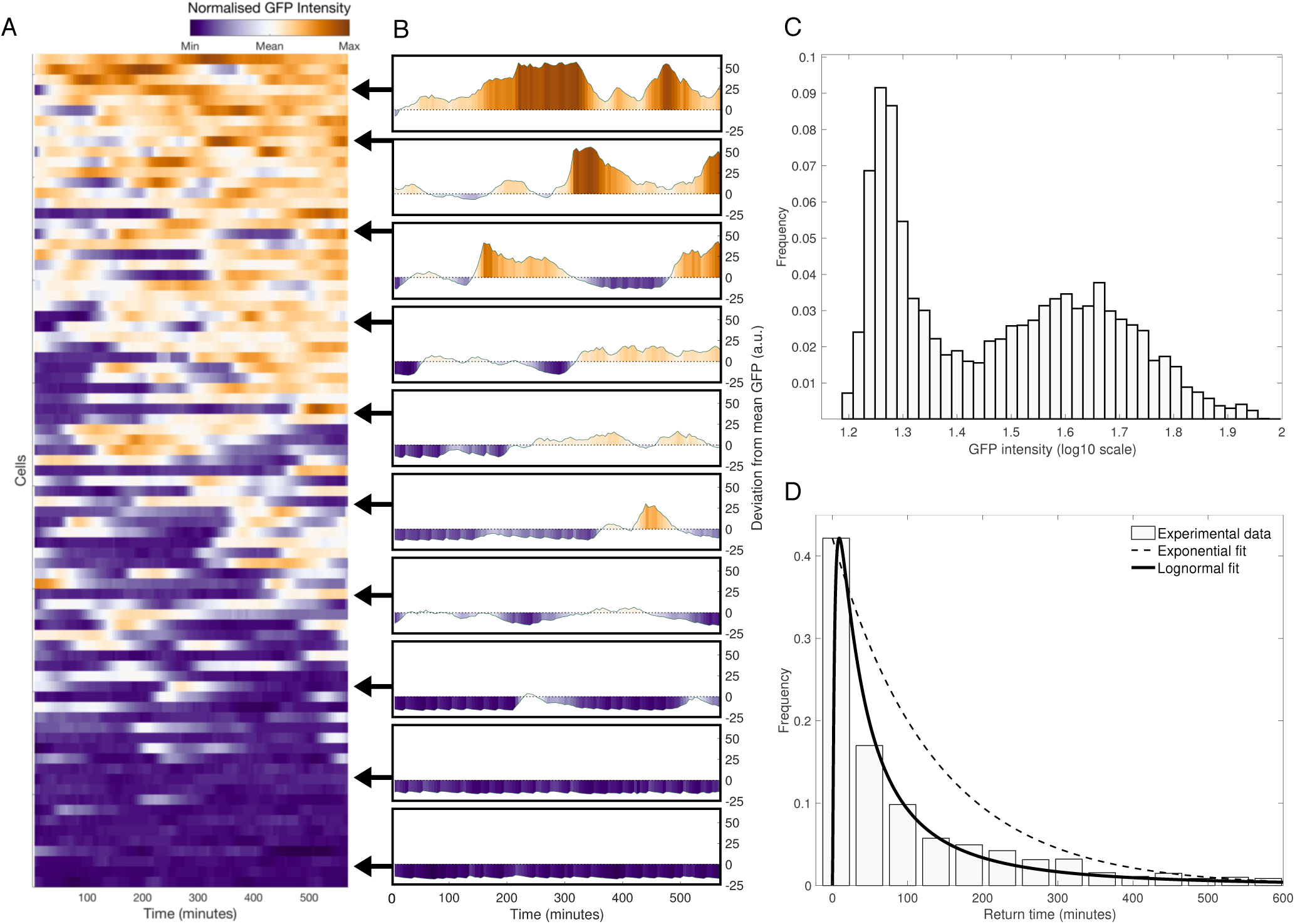
A) Heatmap representing the time-series of fluorescent intensity of 82 cells sorted according to mean fluorescent intensity. Maximum GFP intensity is illustrated with orange, population-level mean GFP is represented with light grey and purple denotes no GFP detected. B) Deviation from the mean fluorescent intensity of a selection of cells. Note that some cells are in the *fliC*-OFF state for the duration of the experiment (bottom), but those cells that are exhibiting fluorescence (top) have a profile of gene expression characterised by non-periodic oscillations. C) Population-level histogram of *fliC* expression obtained from single-cell microscopy observations is bimodal. D) Return time distribution. The dotted and solid lines represent the best-fit lognormal and exponential distributions respectively.

Figure 3D shows an empirical distribution of return times between ON and OFF states as estimated from experimental time-series. One of the defining statistical features of stochastic toggle-switch models is that return times follow an exponential probability distribution: the probability of returning to a basin of attraction is proportional to the time elapsed since escaping.^51, 52^ However, and interestingly, our experimental results do not fit this expectation: in our single-cell data, the time elapsed between ON-OFF switches are not drawn from an exponential distribution (Anderson-Darling test, *p <* 1 × 10^−6^).

Some experimental time-series have a limitation whereby transitions into one state are not followed by a return to the state at the beginning of the observation period. To overcome this, we produced synthetic time-series by iterating a hidden Markov model with the probability of transition between states derived from our gene expression data (see Methods for a description of our quantisation algorithm and Figure S3 for a visual representation of the resulting matrix of transition probabilities). This process shows that the return time distribution between *fliC*-ON and *fliC*-OFF is also not best described by an exponential distribution (Anderson-Darling test, *p <* 1 × 10^−8^). Taken together, both the observed single-cell gene expression data and the synthetic time-series generated using a Markovian description of that data suggest that the underlying mechanism driving *fliC*-ON-*fliC*-OFF transitions is not a stochastic bistable toggle-switch.

### Delayed regulation produces oscillations in the expression of *fliC*

How is this behaviour possible given the architecture of the underlying gene network? We argue that the answer lies in the timescales of feedback regulation. The two negative feedback loops in the flagellar gene regulatory network act on two different levels: FliZ directly inhibits transcription of *ydiV*,^41^ whereas YdiV inhibits FlhDC post-translationally by producing a Ydiv-FlhDC complex and thus indirectly inhibits *fliAZ* by sequestering its positive regulator.^42^

Crucially, post-translational processes such as the protein-protein interaction that leads to the formation of the YdiV-FlhDC complex are known to occur on different timescales (often sub-second) than translation events (minutes).^53^ The negative regulatory effect thus also works on two different timescales and the transcriptional inhibition by FliZ on *ydiV* is delayed relative to the negative regulation of YdiV on FlhDC which is based on the direct interaction of proteins.

Interestingly, delays have been reported in different genetic systems^54^ and they can lead to oscillatory dynamics. For instance, transcriptional delays in a 3-node inhibition loop produce delay-driven oscillations in p53, a tumour suppressor protein involved in DNA-damage response in eukaryotic cells.^55, 56^ Even a simple single-gene network with direct auto-repression of the gene by its own product can produce oscillations,^57^ a feature observed in the segmental clock oscillator in vertebrate development.^58^

To represent delays specific to transcriptional regulation, we modified our model to account for the fact that the concentration of mRNA depends on the levels of regulatory protein present *τ* units of time before the initiation of transcription and, as a result, transcription of *ydiV* at time *t* is inhibited by the concentration of FliZ at time *t* −*τ*. Similarly, FlhDC has a delayed positive interaction on the expression of *fliAZ*. We can then simulate the regulatory dynamics using a system of delay differential equations (equations are described in the Methods).

Figure S4 shows numerical solutions of the delayed system of ordinary differential equations, with fixed delays but of different durations. For small delays, *τ*, the system approaches the *fliC*-ON stable stationary point in an oscillatory manner, but for larger delays, the dynamical system undergoes a bifurcation that creates dynamical oscillations in the expression of FliAZ. The fact that solutions of time-delay differential equations exhibit oscillatory behaviour is unsurprising from a theoretical perspective; it is well-known in dynamical systems theory that delays induce instabilities that may produce oscillations in systems that only have stable attractors when no delays are present.^59, 60^ In particular, delayed negative feedbacks in gene regulation are known to give rise to oscillations for large delays via a Hopf bifurcation.^61^

Although delays can produce sustained oscillations, there is a range of values for *τ* in our system that exhibit transient oscillations whereby the system converges to a stable stationary point (as illustrated in Figure S4B-C). Similarly, a feature of our experimental data is that some cells exhibit sustained non-periodic oscillations in *fliC* for the full duration of our observation but, in some cases, the oscillations collapse and those cells exhibit low fluorescence intensities for the remainder of the observation period. Our data show there are also cells in the *fliC*-OFF state that undergo a sudden spike in *fliC* expression whereafter the system is driven back to the *fliC*-OFF state. In some cases, however, oscillations are resumed and the system fluctuates between *fliC*-ON and *fliC*-OFF states. Indeed, theoretical studies have shown that the introduction of noise in a delayed system can stabilise oscillatory behaviour in parameter regimes where the deterministic mean-field equations converge towards a stable stationary point.^62, 63^

We therefore extended our delayed regulatory model to account for reaction kinetics of the system being generated by a small number of molecules. Previous approaches to stochastic gene dynamics with delays have used computer simulations,^63–65^ derived a delayed master equation^66^ and used virtual reactions.^67^ Here we follow an approach that implements a modified Gillespie algorithm with transcriptional delays^65^ (see Methods); Figure 4A shows a realisation of the stochastic model with delay (*τ* = 1000 units of time). Note that the expression profile exhibits oscillatory behaviour similar to the single-cell time-series of fluorescence intensity in Figures 2B. Also, analogous to the return time distribution estimated experimentally, Figure 4B illustrates that the transition between ON and OFF states in the stochastic model also follows a log-normal distribution (Anderson-Darling test, p=0.249, *H*_0_: the distribution is log-normal).

**Figure 4.**
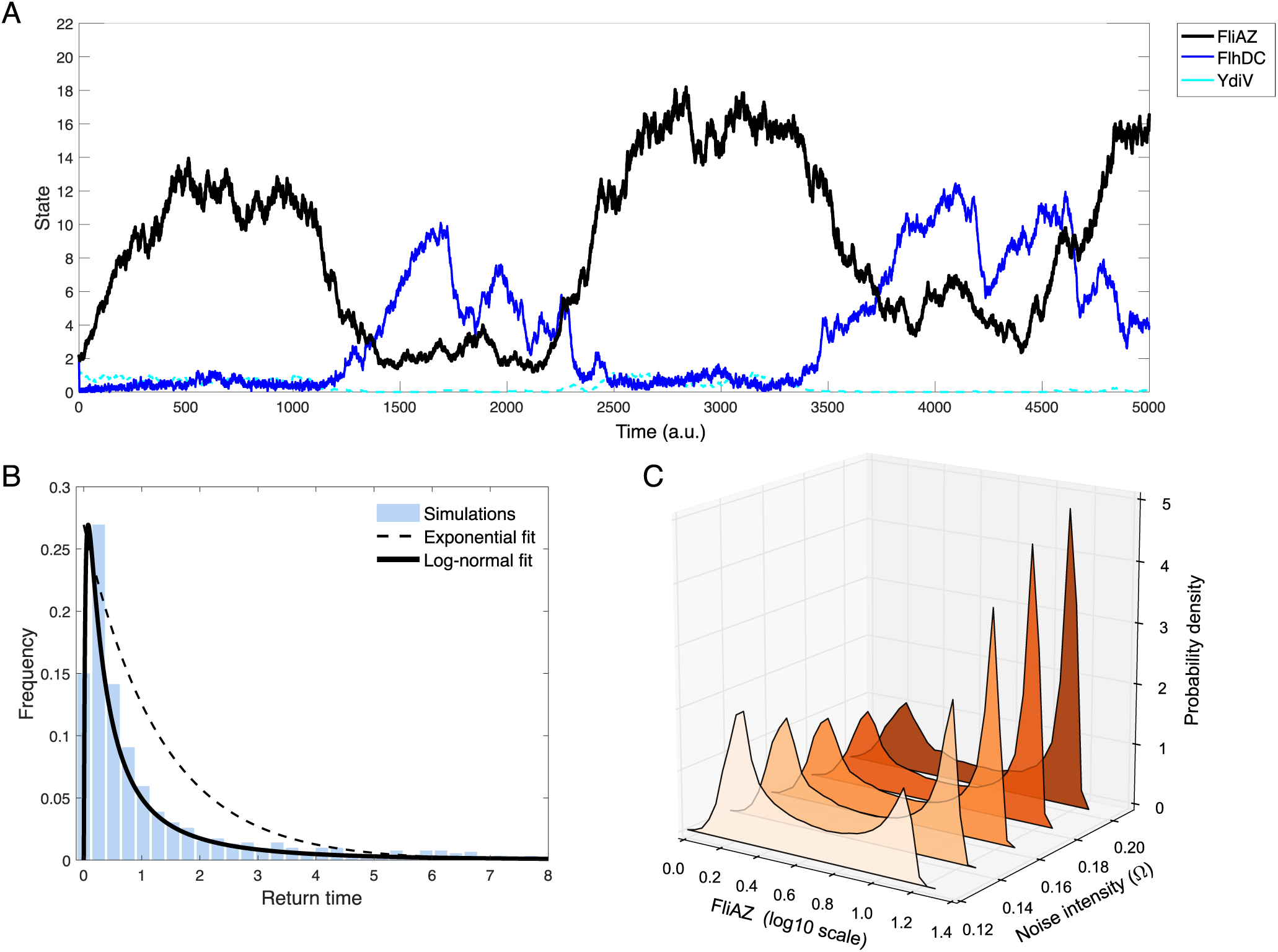
A) Stochastic simulation of the modified Gillespie algorithm to include transcriptional delays (*τ* = 1000, Ω = 0.05). Note how the expression of FliAZ (black solid line) presents oscillatory behaviour. B) Light blue bars represent the return time distribution obtained in our stochastic simulations. As with the experimental data, the return time distribution is best approximated by a log-normal distribution (black solid line) instead of an exponential distribution (black dotted line). C) Frequency distributions of FliAZ expression estimated using different values of Ω. In all cases the distribution is bimodal, but the fraction of the population expressing the virulent phenotype increases at high values of Ω (darker red).

Now, in any physical system with constant volume and temperature, stochastic fluctuations in molecular kinetics decrease when the number of molecules in the system increases. This term is referred to as *system size* and, if we consider it to be constant, then the probability of a reaction ocurring per unit time increases as a function of temperature. To represent this in a stochastic simulation of reaction kinetics, the value associated with the likelihood of each reaction per unit time, a parameter we refer to as *noise intensity* and represent with Ω, should increase. We therefore now use Ω in simulations as a proxy for the environmental temperature in experiments and we can numerically control the intensity of noise in those simulations by implementing different values of Ω.

We then performed stochastic simulations with initial conditions in the *fliC*-OFF state and found, as expected, that the mean escape time from the *fliC*-OFF basin of attraction correlates negatively with Ω (Figure S5C). Indeed, noise-driven escape from the basic of attraction of a (noise-free) stable state happens sooner at greater noise intensity. However, a subsequent prediction of the stochastic-delay model of *fliC* expression dynamics is that the fraction of the population expressing the *fliC*-ON phenotype is also larger when the intensity of noise is higher with less time spent transitioning between either state, features that are reflected in the change of shape of the simulated distribution of FliAZ expression at different values of Ω (Figure 4C).

To test this, we obtained single-cell mean fluorescent intensities in experiments performed at a range of temperatures. Figure 5A shows the deviation from population-level mean fluorescence intensities exhibited by all cells as a stacked dataset. As predicted by the model, we observed a positive correlation between temperature and the fraction of cells in the *fliC*-ON state (Figure 5B). Both the simulations and the single-cell data posses population-level bimodal distributions of fluorescent intensity and, in both cases, observations of the *fliC*-ON mode state are greater at higher noise intensities and temperatures, as can be seen by comparing the frequency distributions shown in Figure 4C and Figure 5C. As a comparison, we performed an analogous procedure using a stochastic model of a bistable, 2-well potential where increasing noise intensity does not mediate the ON-OFF phenotypes like this but instead forces cellular dynamics to spend more time away from each state (Figures S6 and S7).

**Figure 5.**
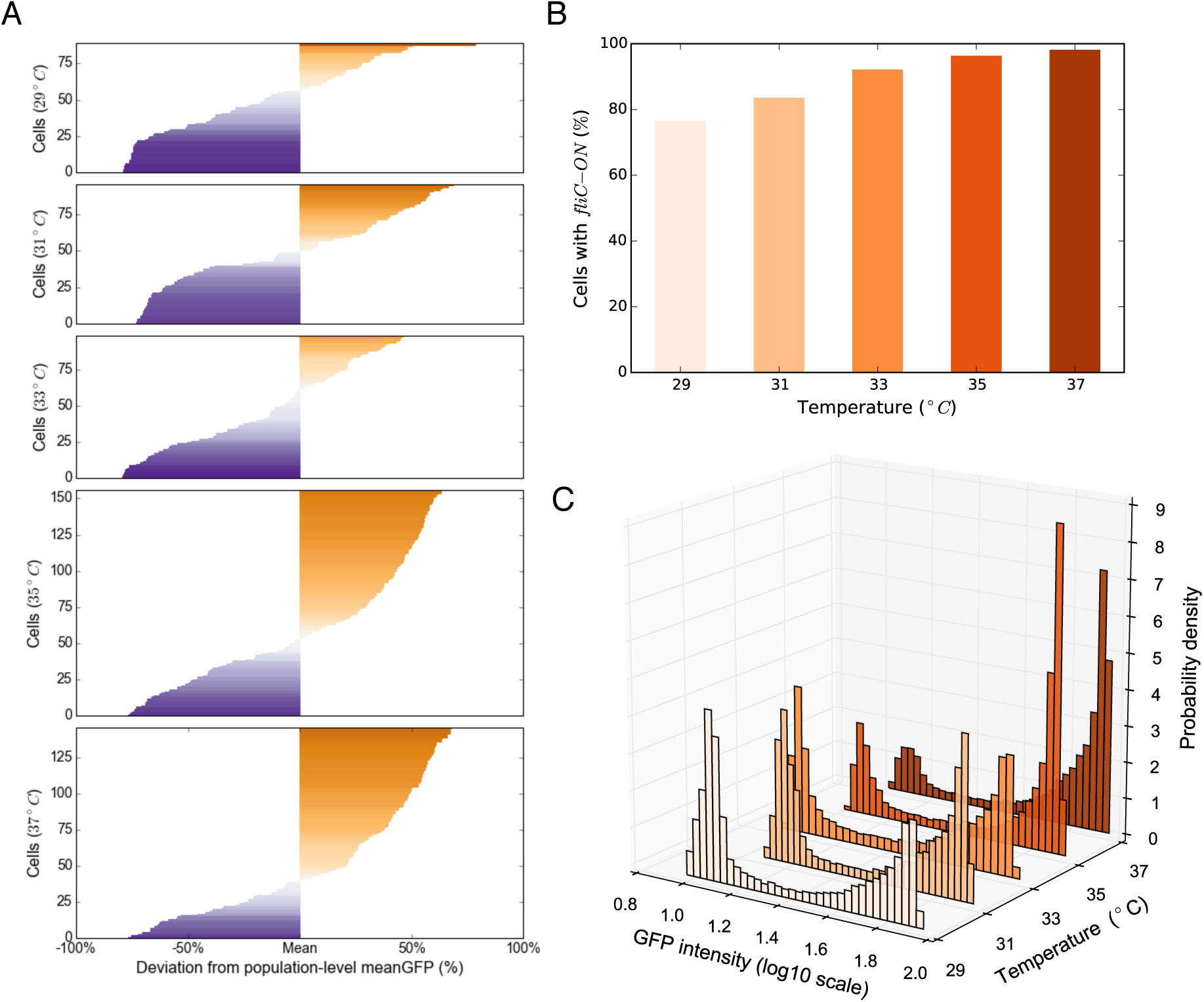
A) Deviation from population-level mean fluorescence of different cells as a function of temperature. Each row corresponds to a single cell, with cells sorted by mean fluorescent intensity. As the temperature increases, so does the fraction of cells expressing mean fluorescence above population-level mean. B) Fraction of cells in the *fliC*-ON basin of attraction as a function of environmental temperature. Notably, at 37*°C*, most of the cells present high levels of *fliC* expression. C) Histogram of GFP expression obtained experimentally. In all cases we obtain bimodal distributions and, as temperature increases, the mode that corresponds to the *fliC*-ON state becomes more prominent.

## Discussion

Although the prevalence of phenotypic heterogeneity is well established^17, 68^ and several benefits of this phenomenon have been described,^1^ the underlying molecular mechanisms that support non-genetic individuality are not well understood. Here we have shown that phenotypic heterogeneity in the expression the flagellar gene *fliC* in *Salmonella* can be explained by dynamically-driven stochastic oscillations. These oscillations result from differences in timescales between the two instances of negative feedbacks in the gene regulatory network: faster post-translational regulation of FlhDC and slower transcriptional regulation of *fliAZ*. Using a computational model that includes both noise and delays, we agree with prior reasoning that different attractors (a.k.a. phenotypes) of the underling dynamical system can be accessed through stochastic fluctuations, but here, by incorporating transcriptional delays, we capture oscillations analogous to the time-resolved, single-cell expression profiles observed experimentally. As a consequence of these oscillations, cells switch between different states more reliably than would be expected through noise-dominated transitions and moreover, those states and the paths between them remain coherent as temperature increases, despite stochastic fluctuations being more likely to dominate a dynamical signal at higher temperatures.

We have shown that multiple gene regulatory dynamical processes can produce analogous population-level bimodal phenotypic distributions, but with very different underling transient dynamics within single cells. Interestingly, both the dynamics discussed in this paper, the noise-driven toggle switch and delay-driven oscillations, are derived from the same network architecture. We argue, therefore, that inferences should not be made about gene regulatory dynamics from population-level observations, instead a bottom-up approach is preferable that employs microfluidic technology and fluorescence microscopy to track the dynamics of gene expression at the single-cell level with high temporal resolution.

In conclusion, heterogeneities in gene expression do not have to be due to irreducible noise in the gene expression apparatus, such as stochasticity imposed by small numbers of molecules, and they can be an inherent feature of a gene regulatory network. We propose that such a hard-wiring of oscillations, and potentially even more exotic dynamics besides, as produced deterministically by the underlying gene regulatory dynamics should have strong implications in terms of the controllability, adaptability, and robustness of a cell’s phenotype. Finally, understanding the functional benefits that oscillatory dynamics bestow on the cells that harbour them represents an important open question.

## Methods

### Bacterial Strains

The bacterial strain used for all experiments in this study is derived from the flagella mutant X8602.^69^ This strain is a Δ*fliC* Δ*fljB* mutant derived from *S.* Typhimurium SL1344. This mutant was used to reduce motility of bacterial cells for microscopy experiments, to avoid loss of cells from the channels. In addition, the cells were carrying the plasmid 01_24.^29^ This plasmid is a derivative of pM967, a low copy number promoter-less plasmid containing *gfpmut2*, as described in.^70^ It is itself derived from pWKS30.^71^ The promoter sequence upstream of *gfpmut2* in 01_24 contains the 233 nucleotides upstream of *fliC* on the *S.* Typhimurium genome.

### Culture Conditions

For microscopy experiments, cells were grown over night in culture tubes (100 mm × 16 mm PP reaction tube, Sarstedt, Numbrecht, Germany) in 5ml LB Lennox (Sigma) medium containing 100*µg/ml* ampicillin (AppliChem). The tubes were incubated in an air incubator at the specified temperatures and shaken at 220rpm. The next morning, cells were diluted 1:100 in 5ml fresh LB Lennox medium containing 100*µg/ml* ampicillin, and incubated for 2-3h to be in exponential phase. Cells were then concentrated approximately 100x by centrifugation (13,000 × g, 2min) and transferred into microfluidic devices for microscopy. For the medium in microscopy experiments, LB Lennox was supplemented with BSA (Sigma), at 150*µg/ml*, and salmon sperm DNA (Sigma), at 50*µg/ml*. These supplements avoid sticking of the cells to PDMS. The temperature during microscopy experiments was controlled by placing the whole microscopy setup in an incubated box (Life Imaging Services, Reinach, Switzerland).

### Microfluidics

Microfluidic devices were fabricated, and experiments performed, as described in Arnoldini et al., 2014,^72^ adapted from a design commonly known as the Mother Machine.^50^ In short, PDMS was mixed with a curing agent (Sylgard 184 Silicone Elastomer Kit, Dow Corning), poured over the silicone wafers, degassed, and cured over night at 80*°*C. Inlet and outlet holes were punched using modified 18G needles, and PDMS chips were bound to glass coverslips (Menzel-Glaser, Braunschweig, Germany) using a UV-Ozone cleaner. Before loading cells, channels were rinsed with culture medium, and cells were then loaded using a pipette. Medium flow was 2*ml/h*, using syringe pumps (NE-300, NewEra Pump Systems) with 60*ml* syringes (IMI, Montegrotte Terme, Italy). Microbore Tygon (S54HL, ID 0.76 mm, OD 2.29 mm, Fisher Scientific) and Teflon (ID 0.3 mm, OD 0.76 mm, Fisher Scientific) tubing was used.

### Microscopy and Image Processing

An Olympus IX81 inverted microscope system with an automated stage, shutters, and hardware autofocus was used. Several different positions were monitored in parallel on the same device, and phase contrast and fluorescence images were taken at every position at 5 min intervals. Images were acquired using an UPLFLN100xO2PH/1.3 phase contrast oil immersion objective (Olympus) and a cooled CCD camera (Olympus XM10). CellM software (Olympus) was used for image acquisition. The light source for fluorescence imaging was a 120W mercury short arc lamp (Xcite 120PC Q), and GFP was imaged using the U-N21001 filter set (450-490 nm ex/500-550 nm em/ 495 nm dichroic mirror, Chroma). Time-series of gene expression were obtained from time-lapse images analysed using the plugin MMJ^72^ for ImageJ.^73^

### Dynamical Model of Gene Regulation

Let us denote the state of the system as *x*(*t*) = (*m*_*z*_, *p*_*z*_, *m*_*v*_, *p*_*v*_, *m*_*d*_, *p*_*d*_) where *m*_*z*_, *m*_*v*_ and *m*_*d*_ represent the concentration of *mRNA* produced by genes the operons *fliAZ, yDiV* and *flhDC* respectively, and with corresponding time-dependant protein concentrations denoted by *p*_*z*_(*t*), *p*_*v*_(*t*) and *p*_*d*_ (*t*). Furthermore, as the expression of *fliC* is directly regulated by *fliAZ*, we will assume that the expression of both genes is correlated. Now, if we consider that binding of proteins occur fast compared to transcription and translation, then we can approximate the gene network dynamics using the following set of ordinary differential equations:

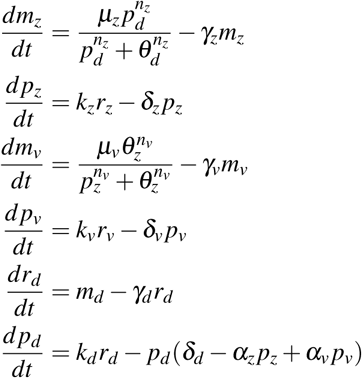

suitably augmented with initial conditions and where *k*_*i*_ denotes translation rate, *γ*_*i*_ mRNA degradation rate, *δ*_*i*_ protein degradation rate and *µ*_*i*_ represents the maximal transcription rate of gene *i. α*_*z*_ and *α*_*v*_ denote the rates of protein-protein interaction of *p*_*d*_ with *p*_*z*_ and *p*_*v*_ respectively. Finally, *θ*_*i*_ and *n*_*i*_ correspond to the expression threshold and cooperativity coefficients of the Hill function used to model the synthesis of mRNA transcribed by gene *i*.

Now, if we consider that the transcriptional inhibition of FliZ on *ydiV* and the activation of *fliAZ* by FlhDC occur with a constant delay *τ* > 0 with respect to the remaining interactions, then we can modify the evolutionary equations for *m*_*z*_ and *m*_*v*_ as follows:

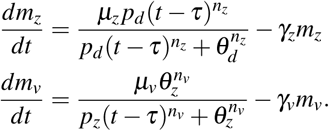

Numerical simulations of the model presented in this paper were performed using standard numerical solvers of Matlab and NumPy with parameter values described in Table S2.

### Computational Simulations with Noise and Delay

We used a Gillespie algorithm to obtain numerical realisations of the stochastic model. Table 1 describes the reactions that underlie the interactions of the three-tier cascade that regulates the synthesis of the flagellum and Figure 1D shows numerical realisations of this algorithm. Gillespie algorithms are based on simulating reactive events and the subsequent changes in the molecular configuration of the system, with propensities of the occurrence of each reaction determined by the reaction rate constant multiplied by a coefficient proportional to the number of reactant molecules in the system, Ω. For the purpose of this paper, we use this parameter as a proxy for temperature. In order to model post-transcriptional delays, we use a modified Gillespie algorithm proposed in.^65^ That is, we assume that some reactions are Markovian with a probability of occurring proportional to the number of substrate molecules and the propensity of the reaction and, using a pseudo-random number generator, we decide the time interval before the next reaction and the next reaction to occur. Delayed reactions are pushed into a stack and performed *τ* units of time later. Simulations of this modified Gillespie algorithm were performed in Matlab.

### Hidden Markov Model

In order to estimate a return time distribution from the *fliC-OFF* basin of attraction, we produced a synthetic time-series of gene expression by iterating a hidden Markov model that underlies a quantised version of our gene expression data. First, we filtered the noise of each experimentally-acquired time-series using a Fast Fourier transform-based low-pass filter applied to the gene expression data of every cell in our experiment. A boolean (*ON-OFF*) quantisation produced a poor description of the expression time-series, so we identified the *n* dominant quantised states from a histogram of states observed at all times, where *n* represents the number of states for the quantisation, and we associated gene expression values to each state. A numerical optimisation algorithm based on a minimal information criterion with *n* as the optimisation parameter showed that *n* = 7 was the number of states that best approximated our data (see Figure S3A for an example). We then used the Causal State Splitting Reconstruction algorithm (CSSR)^74^ to estimate a *n*× *n* matrix of transition probabilities between the quantised states and their quantised between-state trajectories. Finally, by initialising the system into any one of these discrete states and iterating the resulting Markovian model, we could produce synthetic timeseries data with the same between-state transition probabilities as the single-cell data obtained experimentally.

## Acknowledgements

REB was funded during this work by an MRC Discipline Hopping Fellowship G0802611. REB and RPM were supported by EPSRC grant EP/I00503X/1 awarded to REB. RPM was also supported by an IDEA League fellowship, by UNAM-DGAPA-PAPIIT (grant IN209419) and CONACYT Ciencia Básica (grant A1-S-32164). MAr and MAc were funded by grants 31003A 130735 and 31003A 149267 by the Swiss National Science Foundation to MAc. MAr was also supported by a fellowship from the Swiss National Science Foundation (P300PA 160965) and was a nonstipendiary EMBO Fellow (ALTF 344-2015) during this work.

## Supplementary Figures

**Figure S1.**
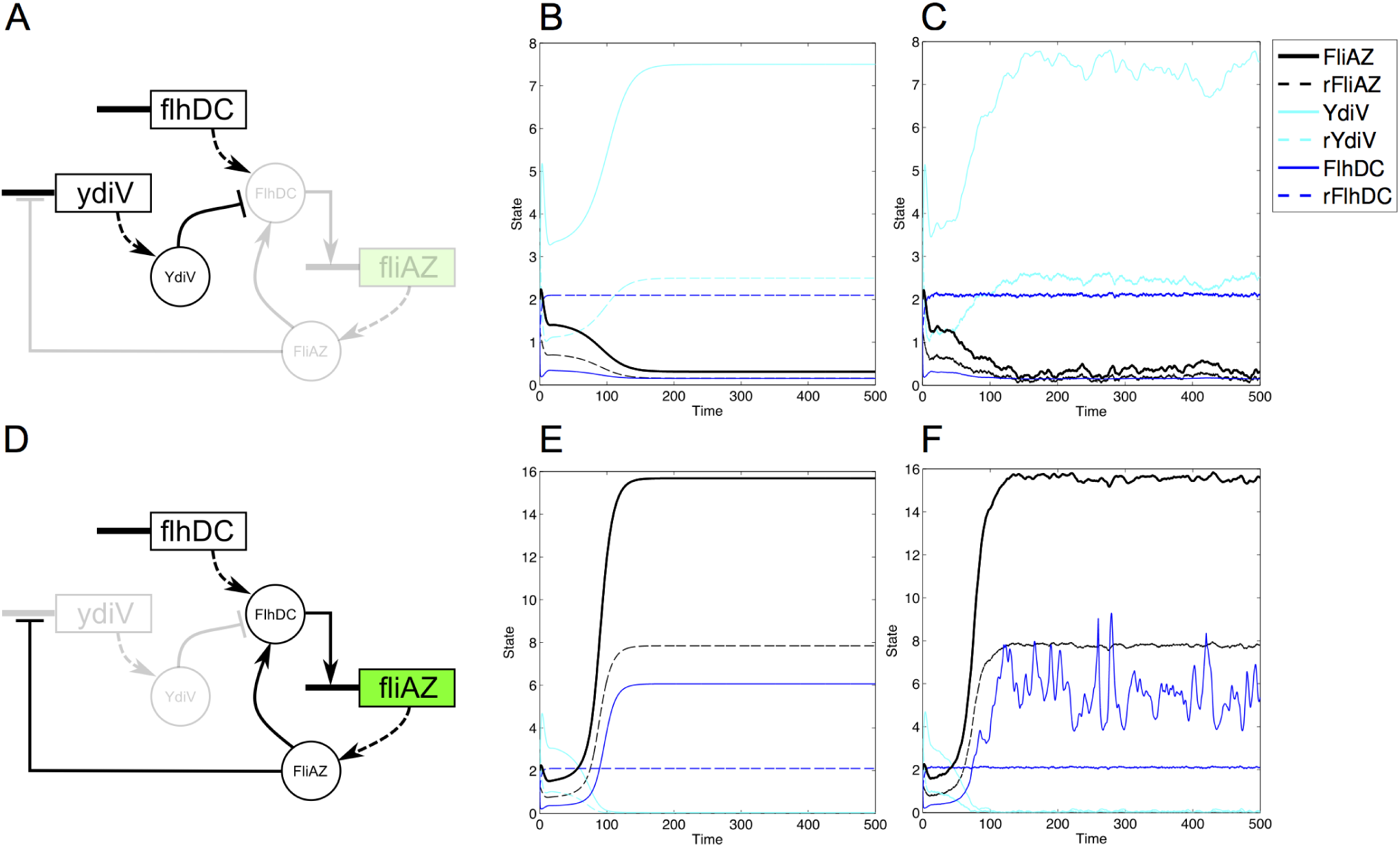
The dynamics presented by the gene regulatory model is bistable. A) If *ydiV* is expressed, then *fliAZ* expression is repressed by post-translationally inactivating its FlhDC. B) Numerical solutions of the deterministic model shows how the expression of *fliAZ* is repressed. Stochastic simulation of the system using the same initial conditions and parameters as in B). D) If *ydiV* is repressed, then the master regulator *flhDC* activates the expression of *fliAZ* and the state of the system is in the *fliC*-ON state. E) and F) Deterministic and stochastic solutions of the model show how in this case the level of expression of *fliAZ* (black solid line) is high.

**Figure S2.**
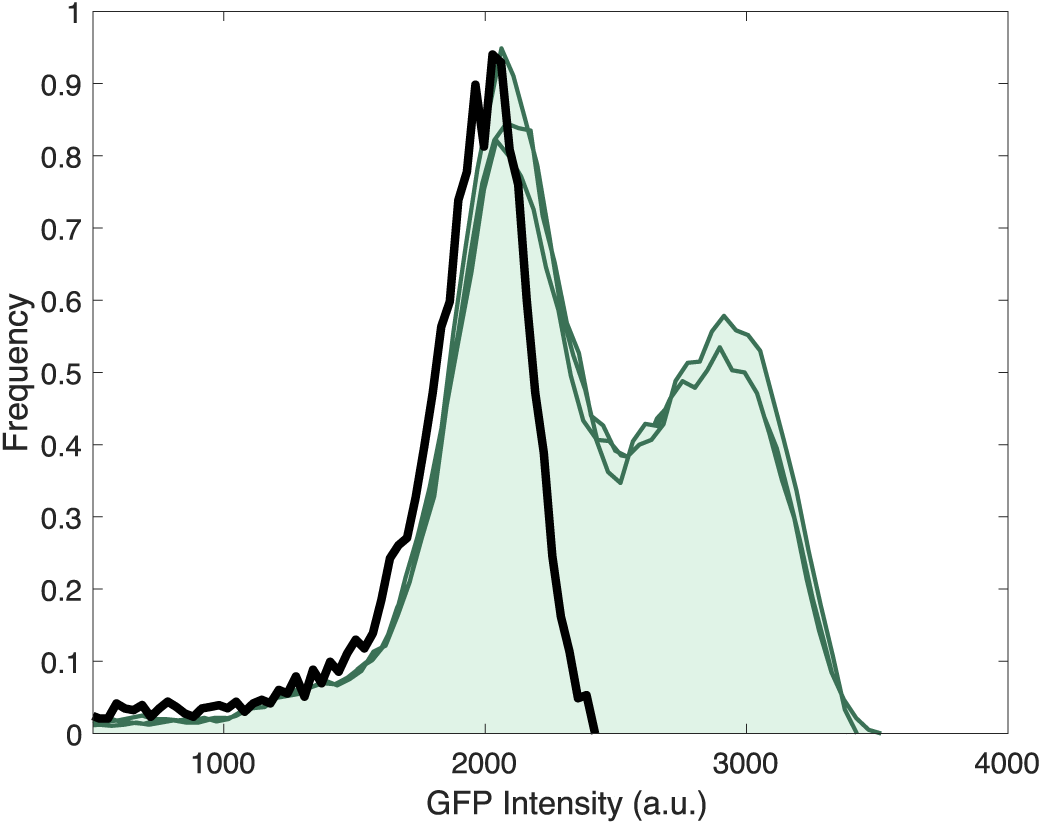
Flow cytometry data showing the population-level fluorescent distribution of a clonal community of Salmonella with a GFP marker under the control of the *fliC* promoter; green lines represent three replicates with a fluorescently-tagged *fliC* strain, while the black line is a non-fluorescent control. Note that the fluorescence histograms of the GFP-tagged cells present a bimodal distribution.

**Figure S3.**
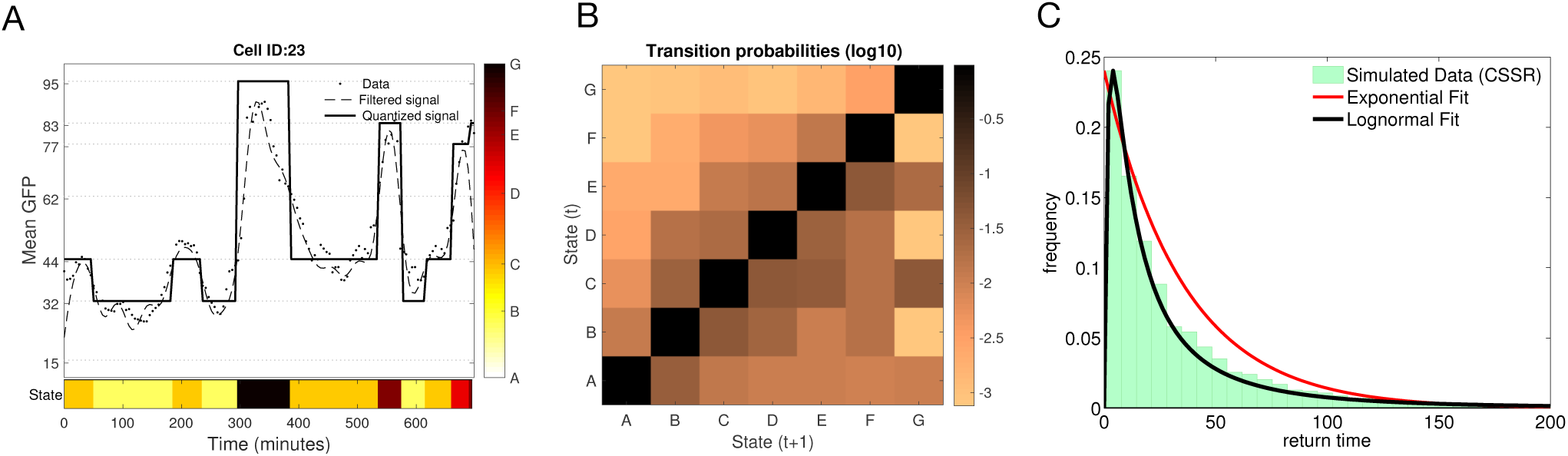
A) Example of a 7-state quantised time-series of GFP intensity (dots represent experimental data, dashed line the filtered signal and the black solid line the discretised time-series). The discrete expression states (represented by letters A-G) of this time-series are colour-coded in the bottom of the figure. B) Matrix of transition probabilities between expression states obtained using a Casual State Splitting Reconstruction algorithm (darker colour denotes a higher transition probability). C) By iterating the transition matrix given an initial state, we produced synthetic data of fliC expression. C) We used these long time-series to estimate a distribution of return times.

**Figure S4.**
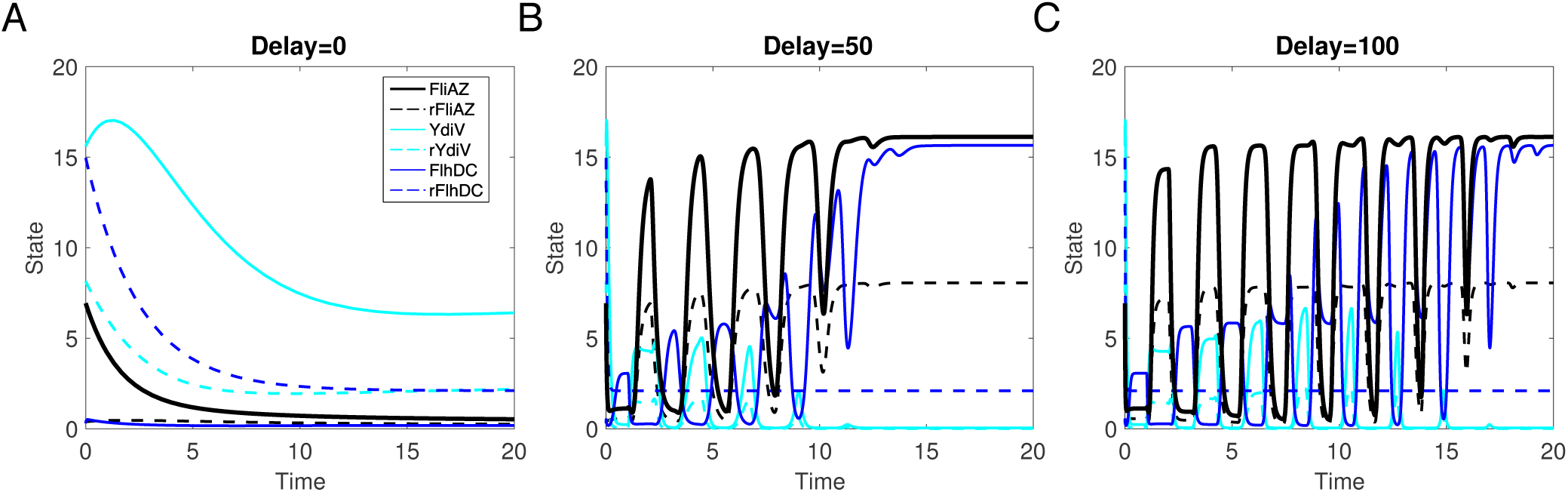
Numerical solutions of the deterministic model for a fixed time-delay, *τ*, performed using the solver *dde* in Matlab. A) If *τ*=0 the system converges to the *fliC*-OFF state. B) At intermediate values of *τ* the system undergoes a bifurcation and starts oscillating. C) For large delays the system converges to an oscillatory regime that dynamically alternates between *fliC*-ON and *fliC*-OFF states until eventually the oscillations collapse and the system converges to a stable stationary point.

**Figure S5.**
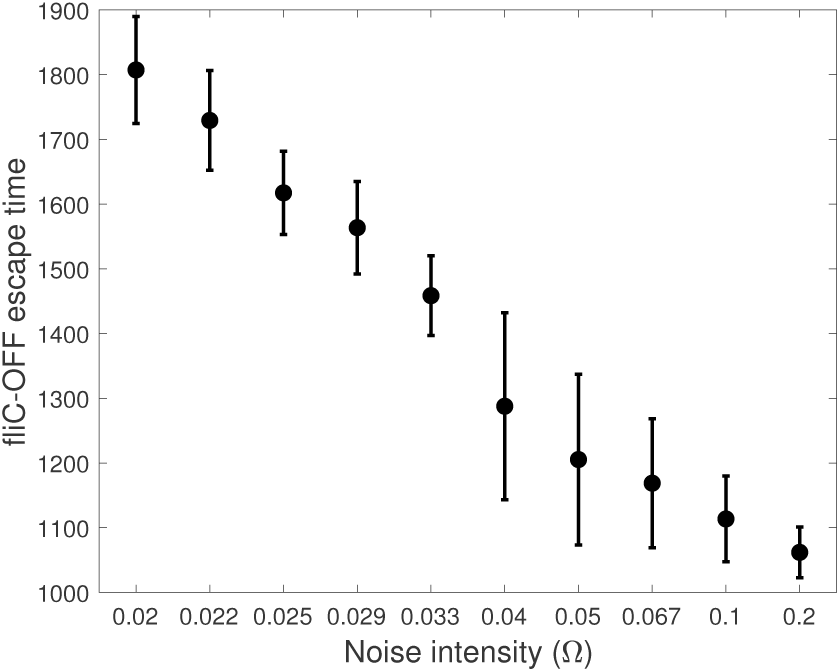
Theoretical mean escape times from the *fliC*-OFF basin of attraction are negatively correlated with the noise intensity: at high values of Ω, the probability of transitioning to the *fliC*-ON basin of attraction is increased.

**Figure S6.**
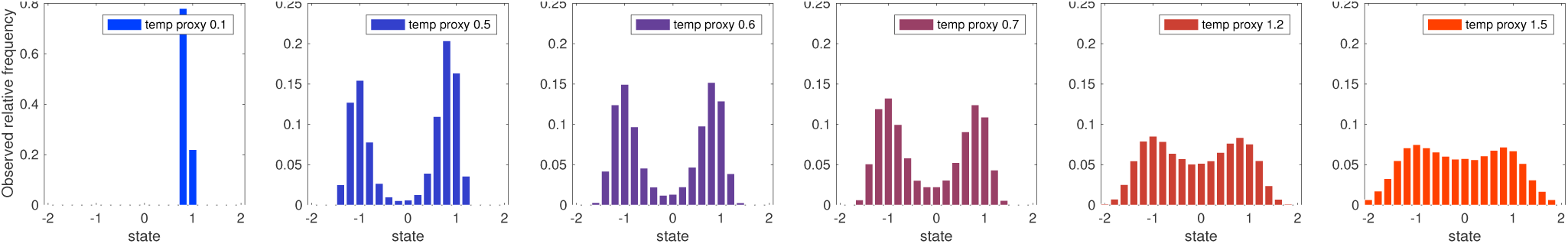
These are histograms of observed states that result when increasing noise intensity in a stochastic model based on a mathematical model of phenotypic ‘particles’ moving in a bistable, 2-well potential: note, because the time states spend close to each phenotype (at *x* = +1 and *x* = −1) reduces as noise increases (ordered from left to right) the system spends more time in intermediate states (near zero *x* = −0). Finally, as noise increases further (the rightmost plot) the histogram approaches a near-uniform distribution wherein the two phenotypes can no longer be distinguished in the data. These data should be contrasted against the experimentally observed outcomes from single-cell observations in Figure 5C that do not have this form. Simulations for this supplementary figure use the stochastic oscillator *d*^2^*x/dt*^2^ + *V*^*′*^(*x*) = *δdx/dt* + *σdW* where *x*(*t*) is said to be the phenotype or state, additive noise intensity *σ* is the temperature proxy, *V* (·) is a 2-well potential, dissipation is *δ* > 0 and *W* is Brownian motion (Matlab codes are supplied). Also see Figure S7 below for more details.

**Figure S7.**
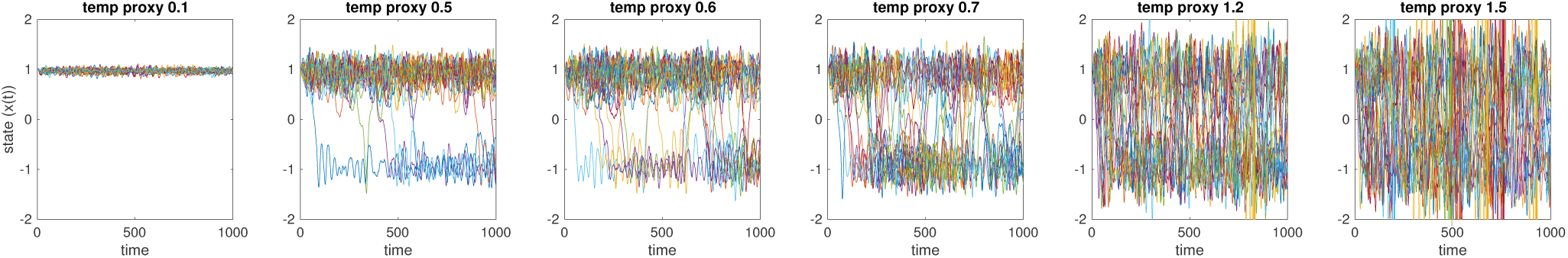
Exemplar stochastic trajectories (*x*(*t*)) realised from numerical solutions of *d*^2^*x/dt*^2^ + *V*^*′*^(*x*) = *δdx/dt* + *σdW* that form the basis of the histograms in Figure S6. They do not resemble the empirical *fliC* trajectories in Figure 3 and the main text explains that the statistics of return times between states is one way of demonstrating this lack of resemblance formally.

**Table S1.** Reactions and propensities of the flagellar gene regulation stochastic model.

| Number | Reaction | Propensity |
| --- | --- | --- |
| 1 | $\rightarrow r_z$ | $w_1 = \frac{m_z D^{n_z}}{D^{n_z} + \theta_d^{n_z}}$ |
| 2 | $\rightarrow r_v$ | $w_2 = \frac{m_v \theta_z^{n_v}}{Z^{n_v} + \theta_z^{n_v}}$ |
| 3 | $\rightarrow r_d$ | $w_3 = 0$ |
| 4 | $r_z \rightarrow$ | $w_4 = \gamma_z r_z$ |
| 5 | $r_v \rightarrow$ | $w_5 = \gamma_v r_v$ |
| 6 | $r_d \rightarrow$ | $w_6 = \gamma_d r_d$ |
| 7 | $r_z \rightarrow Z$ | $w_7 = k_z r_z$ |
| 8 | $r_v \rightarrow V$ | $w_8 = k_v r_v$ |
| 9 | $r_d \rightarrow D$ | $w_9 = k_d r_d$ |
| 10 | $Z \rightarrow$ | $w_{10} = \delta_z Z$ |
| 11 | $V \rightarrow$ | $w_{11} = \delta_v V$ |
| 12 | $D \rightarrow$ | $w_{12} = \delta_d D$ |
| 13 | $ZD \rightarrow D$ | $w_{13} = \alpha_z ZD$ |
| 14 | $VD \rightarrow$ | $w_{14} = \alpha_v VD$ |

**Table S2.** Parameters used in the numerical solutions of the deterministic model

| *Parameter* | *Value* | *Description* |
| --- | --- | --- |
| $k_i$ | (1.2, 0.9, 1.4) | translation rate of gene $i$ ($z, v, d$) |
| $\gamma_i$ | (0.1, 0.3, 0.4) | mRNA degradation rate ($z, v, d$) |
| $\delta_i$ | (0.6, 0.3, 0.4) | protein degradation rate ($z, v, d$) |
| $\mu_i$ | (0.81, 0.8, 0.84) | maximal transcription rate ($z, v, d$) |
| $\theta_i$ | (1.1, 1.2) | Hill expression threshold ($z, v$) |
| $n_i$ | 2 | Hill cooperativity coefficient |
| $\alpha_z$ | 0.02 | $p_d$ and $p_z$ protein interaction rate |
| $\alpha_v$ | 2.5 | $p_d$ and $p_v$ protein interaction rate |

